# Paeniclostridium sordellii NanS works synergistically with TcsL to increase cytotoxicity and the severity of pathogenesis in the murine uterus

**DOI:** 10.1101/2023.06.22.546043

**Authors:** Sarah C. Bernard, Heather K. Kroh, M. Kay Washington, D. Borden Lacy

## Abstract

*Paeniclostridium sordellii*, a pathogen capable of causing lethal uterine infections post-partum or post-abortion, produces a variety of potential virulence factors. Lethal toxin (TcsL) has been found to be essential for lethal *P. sordellii* infections, but the physiological role of NanS, a sialidase, remains elusive. Here, we purify enzymatically active *P. sordellii* NanS and report the crystal structure. We show that NanS works in a synergistic manner with TcsL to increase cytotoxicity and cell rounding of tissue culture cells. Using a mouse model of hormone-dependent uterine intoxication, we show that NanS augments TcsL-induced lethality in diestrus mice. In estrus mice, the combination of NanS and TcsL significantly increases uterine histologic damage compared to challenge with TcsL alone. This suggests that NanS enhancement of TcsL-induced uterine epithelial injury may potentiate TcsL access to the tissue when mucus is present.

**Author Summary:** *P. sordellii* produces several proteins predicted to contribute to lethal uterine infections following childbirth, stillbirth, or abortion. The role of lethal toxin (TcsL) in causing lethal outcomes, has been well documented, but little is known of how NanS, a sialidase, contributes to disease development. Previous reports have suggested a role for NanS in increasing the tissue culture cell toxicity from *P. sordellii* supernatants. In this study, we show in cell culture and using a uterine mouse model that TcsL and NanS work together to increase cell cytotoxicity and to worsen disease outcome. This finding suggests an accessory role for NanS to enhance *P. sordellii* pathogenesis when host environmental conditions reduce the influence of TcsL.

## Introduction

*Paeniclostridium sordellii* is a human pathogen capable of causing lethal intrauterine infections following childbirth or abortion. *P. sordellii* produces two major exotoxins, TcsL and TcsH, as well as additional virulence factors, such as a phospholipase, a hemolysin/cytolysin, and a sialidase, NanS. Only TcsL has been found to be essential for lethal *P. sordellii* infections (PSI) [1]. Few studies, however, have been done to show how these other virulence factors participate in pathogenesis.

Bacterial sialidases are responsible for cleaving terminal sialic acids from host glycoproteins and glycolipids and are found to have roles in bacterial metabolism [2–4], biofilm development [5], bacterial adherence [6], and in increasing cytotoxicity of individual toxins [6–10]. In the case of NanS, it has been reported to increase the proliferation of HL-60 cells *in vitro*, suggesting that it may play a role in the characteristic leukemoid reaction (LR) in PSI [11]. However, a subsequent study infected animals with a *P. sordellii nanS*-knockout that still resulted in the developed of a LR, suggesting other virulence factors were responsible [12]. Additionally, NanS has been shown to work synergistically with an unidentified toxin to increase the cytotoxicity of *P. sordellii* [13]. Of note, the *nanS* gene is present in all *P. sordellii* isolates [14], suggesting that it may play an important accessory role in PSI.

Features of sialidase active sites are largely conserved, having a glutamic acid–tyrosine nucleophilic charge relay, an aspartic acid acid/base, and the arginine triad [4]. It has been shown that NanS predominantly cleaves sialyl α2-3 linkages over sialyl α2-6 and α2-8 linkages [15]. In addition to recognition of sialyl linkages, sialidase cleavage is determined by recognition of the carbohydrate structures underlying the sialic acids as well as the overall structure of the glycoprotein or glycolipid [16].

The luminal surface of the uterine epithelium is lined with a carbohydrate-rich layer known as the glycocalyx. Many of the glycoproteins of the glycocalyx network are capped with sialic acids [17], providing an overall negative charge to the mammalian cell surface and creating electrostatic repulsion [18]. These terminal sialic acids also contribute to the overall structure of the glycocalyx, and as shown by sialidase disruption, are key regulators of micro-vessel permeability [19]. Slight modifications in cell surface glycosylation patterns can lead to drastic changes in cellular behavior [16]. In the case of *C. perfringens*, sialidase removal of surface sialic acids was suggested to reduce the electrostatic repulsion, facilitating the binding of toxins to their receptors [18].

Lining the top of the glycocalyx is a layer of mucus largely consisting of the sialoglycoproteins: mucins and immunoglobulins. Mucus and glycocalyx layers are the first and principal barriers protecting the underlying epithelium from pathogenic invasion. Bacterial sialidases, however, can participate in mucus degradation and immunoglobulin hydrolysis, ultimately leading to increased susceptibility to uterine infections [20,21]. In this study, we sought to elucidate the physiological role of NanS in *P. sordellii* pathogenesis using both *in vitro* and *in vivo* techniques.

## Results

### Crystal structure and activity assay of recombinant NanS

To begin to assess the role of NanS in *P. sordellii* pathogenesis, we prepared a plasmid for recombinant protein expression. Following expression and purification of the recombinant protein, we tested NanS for sialidase activity using 4 mM of the substrate 5-bromo-4-chloro-3-indolyl α-D-N-acetylneuraminic acid as previously described [7]. We found NanS to be catalytically active in a dose-dependent manner (**Fig 1A**).

**Fig 1.**
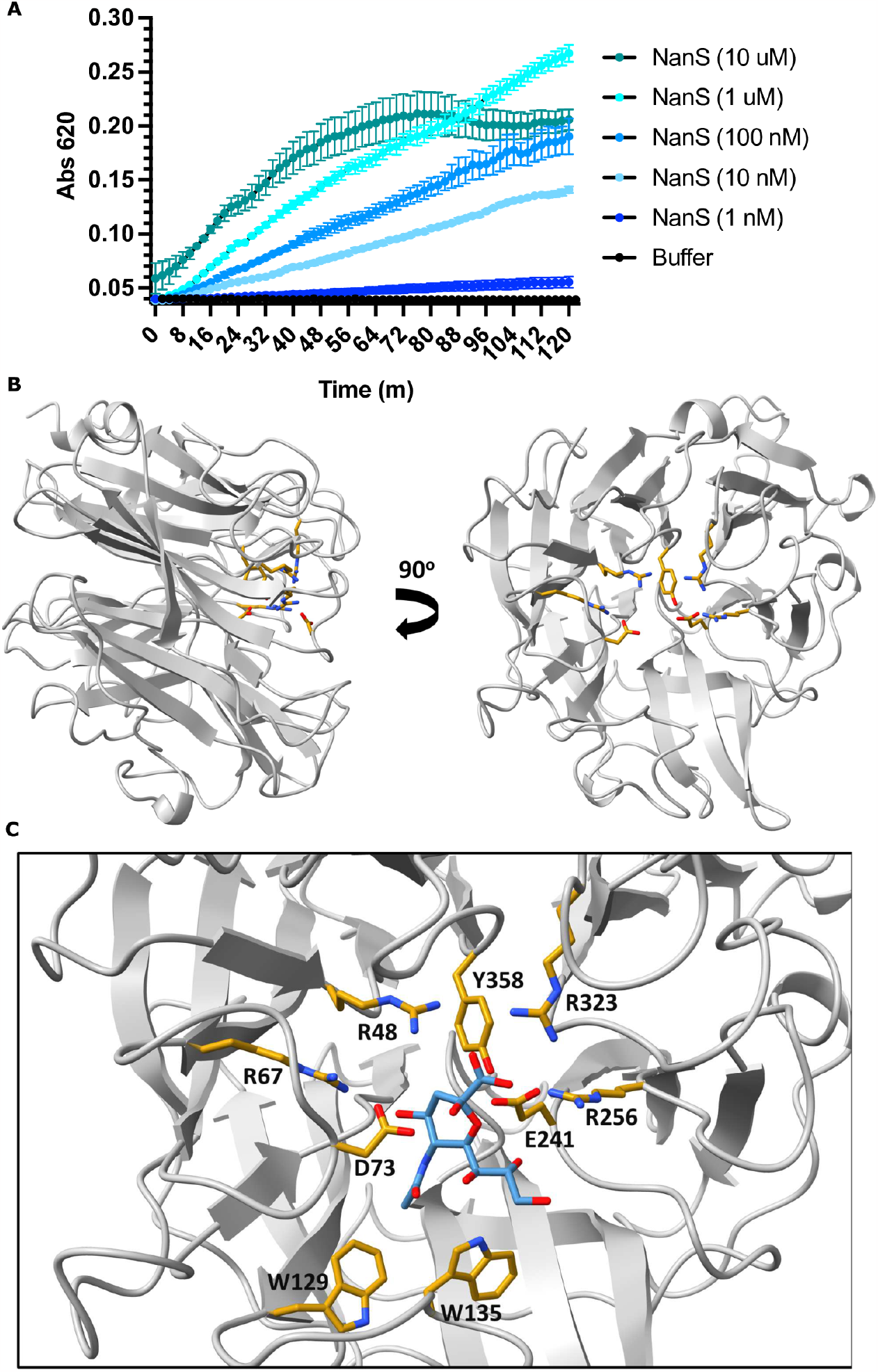
Sialidase activity assay. (**A**) Sialidase activity assay using 4 mM 5-bromo-4-chloro-3-indolyl α-D-N-acetylneuraminic acid of NanS serially diluted 10-fold. (**B**) Crystal structure of NanS with the residues (*yellow*) of the conserved catalytic site with nitrogen (*blue*) and oxygen (*red*) depicted. (**C**) Inset of NanS catalytic site residues modeled with the sialic acid (*light blue*) from *Clostridium perfringens* NanI (PDB 2BF6).

We then performed X-ray crystallography to elucidate the structure of NanS at 1.79-Å resolution (**Table 1**). As with other sialidases, NanS adopts a six-bladed β-propeller fold which contains the catalytic active site (**Fig 1B**) [22,23]. While many bacterial sialidases possess a lectin binding domain inserted ahead of, within, or after the β-propeller fold, NanS appears to only contain the catalytic domain. The active site residues of NanS are highly conserved with those of *C. perfringens* NanI, whose structure was determined alone and in complex with substrate and transition-state analogues [23]. We used the substrate-bound structure of NanI (2BF6) to model sialic acid binding within the NanS active site (**Fig 1C)**. The conservation supports a mechanism where Asp-73 acts as an acid/base proton donor which pushes the C-2 position of the sugar ring into proximity of Tyr-358, which can then serve as a nucleophile. This nucleophile activity is supported by the neighboring Glu-241 which can act as a general base. The sialic acid is oriented in the active site by four arginines, R48, R67, R256, and R323, and a hydrophobic pocket, which in the case of NanS appears to be supported by Trp-129 and Trp-135. Based on these conserved features, we predict that NanS is acting as a hydrolytic sialidase with a two-step double-displacement mechanism that retains the configuration of the *α*-configuration of the glycosidic bond [4].

**Table 1.** Crystallographic data collection and refinement statistics

| Protein | *P. sordellii* NanS |
| --- | --- |
| **Data Collection** |  |
| Wavelength (Å) | 0.97856 |
| Resolution (Å) | 19.77 - 1.79 (1.86 - 1.79) |
| Space group | P2<sub>1</sub>2<sub>1</sub>2<sub>1</sub> |
| Cell dimensions |  |
| *a*, *b*, *c* (Å) | 89.1, 90.9, 129.9 |
| α, β, γ (°) | 90.0, 90.0, 90.0 |
| Total reflections | 482,840 (41,682) |
| Unique reflections | 95,247 (9,656) |
| Multiplicity | 5.1 (4.3) |
| Completeness (%) | 95.7 (98.2) |
| *I*/σ*I* | 16.1 (2.5) |
| Wilson B-factor | 17.19 |
| R<sub>merge</sub> | 0.073 (0.633) |
| R<sub>meas</sub> | 0.082 (0.714) |
| R<sub>pim</sub> | 0.035 (0.321) |
| CC<sub>1/2</sub> | 0.999 (0.773) |
| CC* | 1 (0.934) |
| **Refinement** |  |
| Resolution (Å) | 19.84-1.79 |
| No. of reflections | 95,216 |
| R<sub>work</sub>/R<sub>free</sub> | 0.150/0.183 |
| No. of non-hydrogen atoms |  |
| Protein | 8,502 |
| Water | 951 |
| Total | 9,453 |
| Average B-factor (Å<sup>2</sup>) | 20.88 |
| RMSD values |  |
| Bond lengths (Å<sup>2</sup>) | 0.005 |
| Angles (°) | 0.810 |
| Ramachandran plot (%) |  |
| Most favored | 96.8 |
| Allowed | 2.9 |
| Disallowed | 0.4 |
| Rotamer outliers (%) | 0.0 |

Statistics for the highest-resolution shell are shown in parentheses.

### NanS potentiates TcsL cytotoxicity *in vitro*

In a previous report, NanS was found to work synergistically with an unidentified toxin to increase the cytotoxicity of *P. sordellii* [13]. We wanted to test whether NanS could increase the cytotoxicity of supernatants from various *P. sordellii* strains: ATCC9714, DA108 [24], HH310 [25], and/or ORE2019 [26]. Vero cells were treated with NanS followed by incubation of a ten-fold dilution of *P. sordellii* supernatants grown for 48 hours. ATCC 9714 strain, compared to the others, had an increased cytotoxicity (**S1 Fig A**) and resulted in cell rounding (**S1 Fig B**) following NanS treatment.

One major distinction of ATCC 9714 strain over DA108, HH310, and ORE2019 is the presence of the lethal toxin, TcsL. While the prior report suggested a toxin other than TcsL, we were curious whether NanS would affect the activity of TcsL. We performed *in vitro* cytotoxicity assays where we treated HUVECs with 10 μM NanS for three hours and then assayed at 24 hours for ATP levels as an indicator of viability. NanS alone was not cytotoxic to the cells. However, preincubation of HUVECs with NanS, followed by intoxication of TcsL showed that NanS works synergistically with TcsL to increase cytotoxicity (**Fig 2**). The effect was modest however, and only evident when NanS was added at high concentration (10 μM), leading us to wonder if this synergy would be relevant *in vivo*.

**Fig 2.**
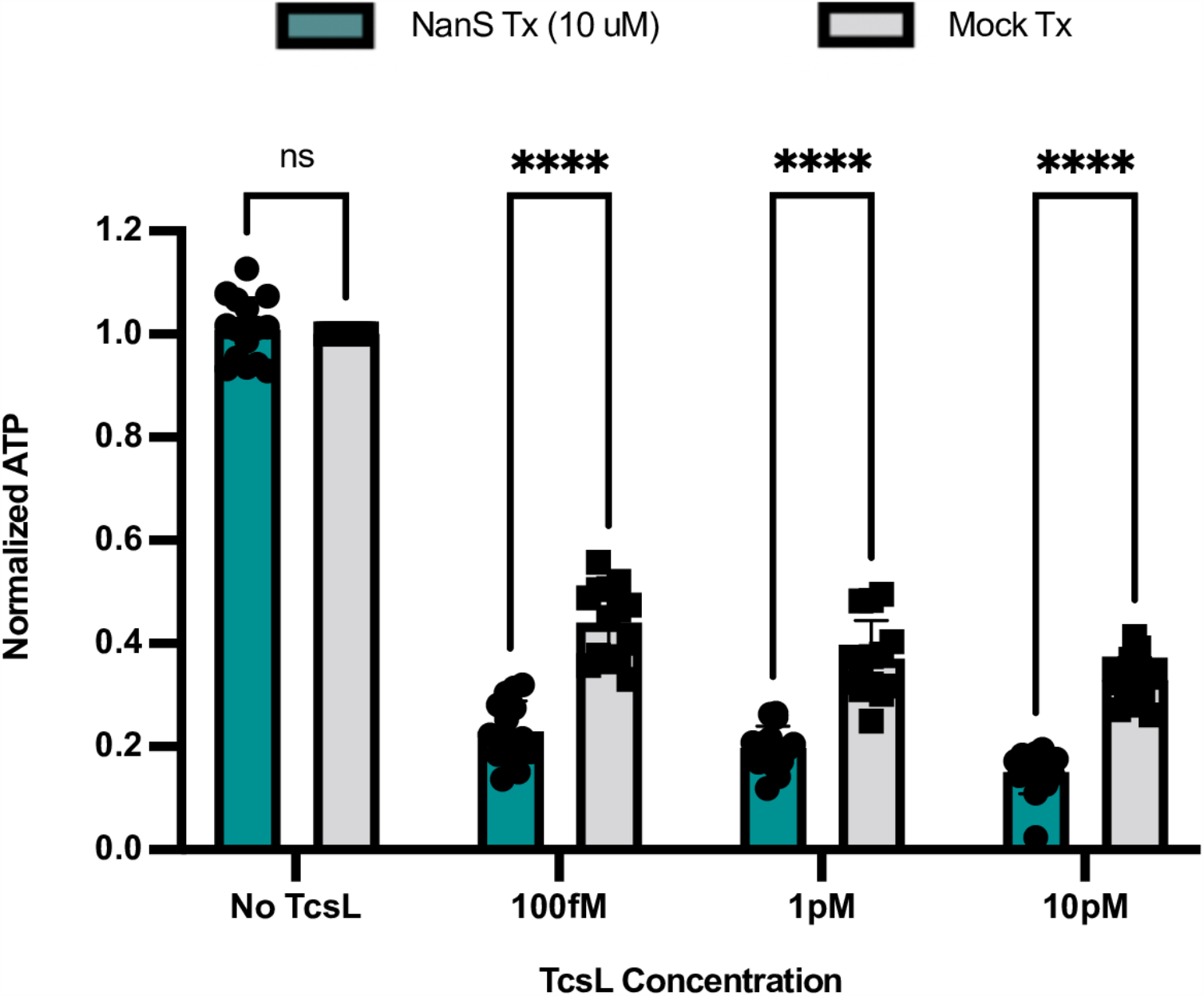
NanS potentiates the cytotoxicity of TcsL. Viability assay of HUVECs treated with 10 μM NanS or mock-treated for 3 hr followed by TcsL intoxication for 24 hr. ATP was measured as a readout of viability and normalized to signal from mock treated cells. Viability data are from four independent experiments with plotting data from technical replicates. Sidak’s multiple comparison (2-way ANOVA) test was used with statistical significance set at a p value of < 0.05.

### NanS increases the rate of Vero cell rounding following TcsL intoxication

It has been reported that the primary target of TcsL during PSI is the vascular endothelium [27]. Therefore, it seemed reasonable for us to do our *in vitro* cell assays with an endothelial cell line such as HUVECs. However, the first cellular barrier TcsL would need to cross in the uterus would be the epithelium and these cells are frequently protected by a layer of mucus. We began to wonder if NanS acted to increase TcsL cytotoxicity on the uterine epithelium, rather than the endothelium.

In our hands, epithelial cells are not very sensitive to TcsL in viability assays, so to test this, a cell rounding assay was performed where NanS-treated or mock-treated Vero-GFP (epithelial) cells were intoxicated with TcsL. Compared to HUVECs, Vero cells were more sensitive to NanS treatment, and less sensitive to TcsL intoxication. For this reason, in contrast to the HUVEC viability assay, we reduced the concentration of NanS to 100 nM, and intoxicated with 100 pM, 10 pM, and 1 pM TcsL. We found that cells pre-treated with 100 nM NanS for 3 hours had a statistically significant increased rate of rounding following TcsL intoxication of 100 pM, 10 pM, and 1 pM, respectively, compared to mock treatment (**Fig 3A-3C**).

**Fig 3.**
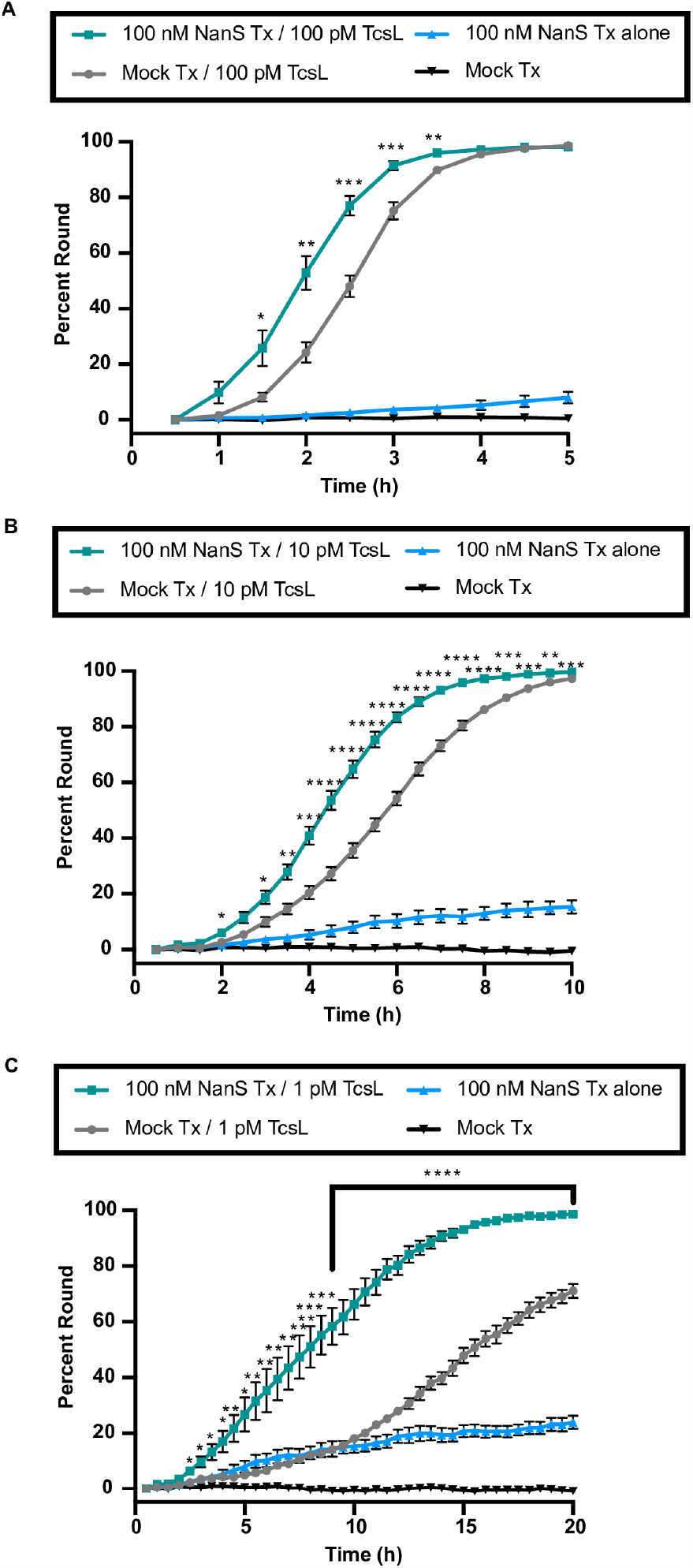
NanS increases the rounding of Vero-GFP cells following TcsL intoxication. Cell rounding percentage from three independent experiments of Vero-GFP cells treated with 100 nM NanS or mock buffer for 3 hr followed by TcsL intoxication at 100 pM (**A**), 10 pM (**B**), or 1 pM (**C)** for 5-20 hr. Data are the accumulation of three independent experiments. Dunnett test for multiple comparisons was used with statistical significance set at a p value of <0.05. (*, P ≤ 0.05; **, P ≤ 0.01; ***, P ≤ 0.001; ****, P ≤ 0.0001). **(A-C)** NanS/TcsL rounding curves were statistically significant from TcsL rounding curves.

### NanS potentiates TcsL-mediated disease in transcervical instillation animal model

We next wanted to assess the importance of NanS in PSI pathogenesis in the context of our transcervical (TC) instillation animal model as previously described [28]. We performed a TC instillation of recombinant NanS and TcsL into animals in diestrus. As shown in Fig 4, 10 ng TcsL transcervically instilled in diestrus mice can kill 50% of mice by day 4. When 1.5 μg NanS was instilled with 10 ng TcsL, all animals were dead before 24 hours, while animals instilled with NanS alone resulted in no signs of sickness or disease (**Fig 4A**). Uterine tissues were harvested upon euthanization and processed for histology. H&E-stained tissue was scored from mild to severe in edema, acute inflammation, and epithelial injury by a pathologist blinded to the experimental conditions (**Fig 4B**). Uterine tissue from control diestrus mice were found to have normal/mild scoring in all criteria, and mice transcervically instilled with 10 ng TcsL or 1.5 μg NanS alone were not found to be statistically significant from control. When NanS was co-administered with 10 ng TcsL, scoring was increased to moderate inflammation and severe epithelial injury with statistical significance. This experiment clearly suggests that NanS can potentiate TcsL *in vivo* and supports a role for NanS on the epithelium.

**Fig 4.**
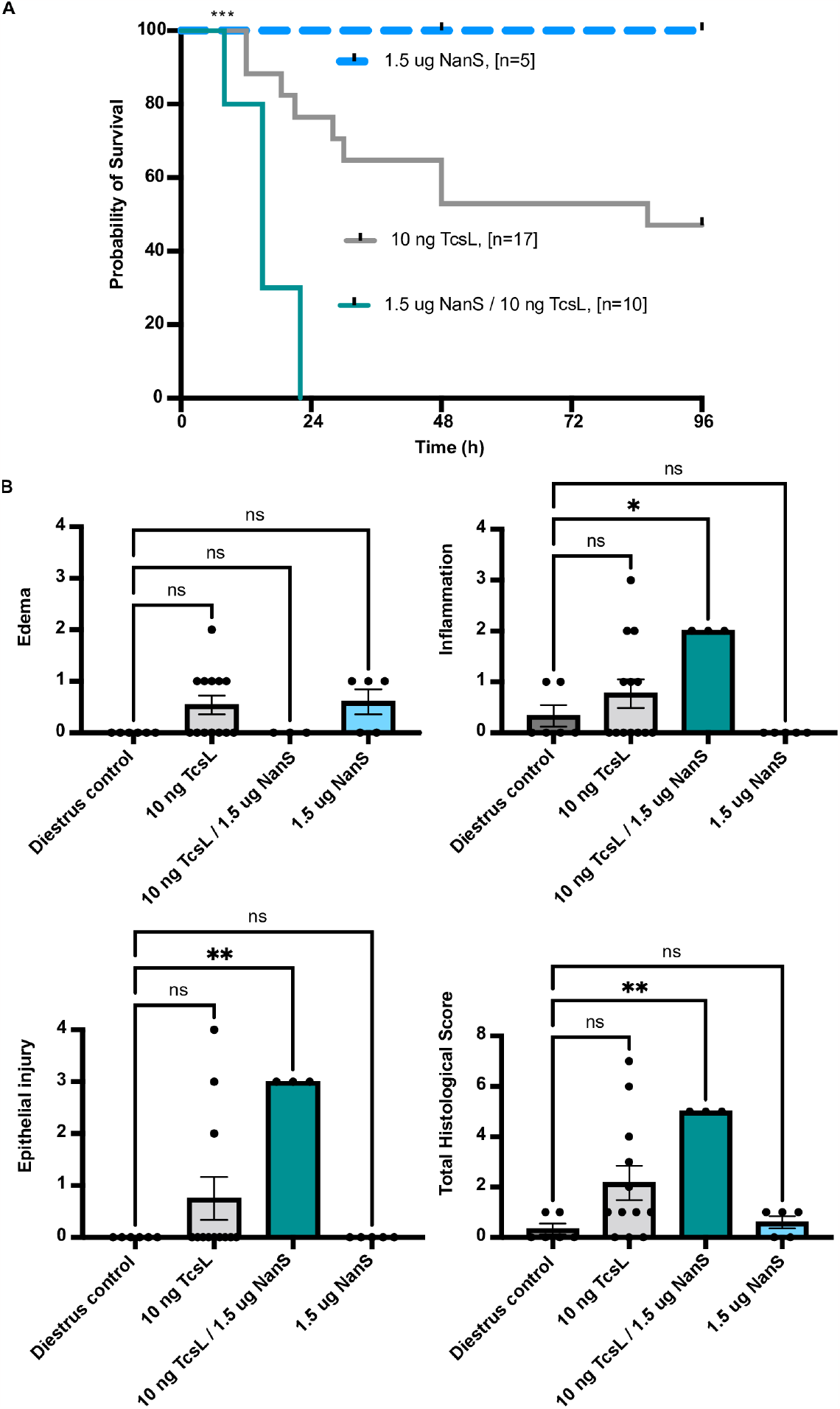
NanS works synergistically with TcsL to exacerbate *P. sordellii* pathogenesis. **(A)** Survival curve of mice that were subcutaneously administered medroxyprogesterone acetate on Day -5 to induce diestrus, followed by transcervical instillation of 1.5 μg NanS, 10 ng TcsL, or 1.5 μg NanS / 10 ng TcsL on Day 0. Animals were weighed/monitored for four days. **(B)** Histological scoring of edema, acute inflammation, and epithelial injury of uterine tissues from moribund mice or at end of study from diestrus mice transcervically instilled with PBS-control, 10 ng TcsL alone, 10 ng TcsL / 1.5 μg NanS, or 1.5 μg NanS alone. Dunnett’s (one-way ANOVA) multiple comparisons test was used with statistical significance set at a p value of <0.05. Some survival data and histological scores from 10 ng TcsL instilled mice have been previously published [28]. Additional representative mice (n=5) were performed in this study and included.

Additionally, we transcervically instilled 20 ng TcsL, 15 μg NanS, or co-instilled 20 ng TcsL and 15 μg NanS in animals in estrus. Animals instilled with TcsL alone or NanS alone survived the study without any signs of disease or sickness. We found, however, that one of ten mice succumbed to co-instillation of TcsL and NanS (**Fig 5A**). Uterine tissues were harvested upon euthanization and processed for histology. H&E-stained tissue was scored following criteria described above (**Fig 5B**). Control estrus mice and mice transcervically instilled with 20 ng TcsL or 15 μg NanS alone were assigned scores of normal/mild in all criteria. When NanS was co-administered with 20 ng TcsL, scoring was increased to moderate inflammation and moderate/severe epithelial injury with statistical significance.

**Fig 5.**
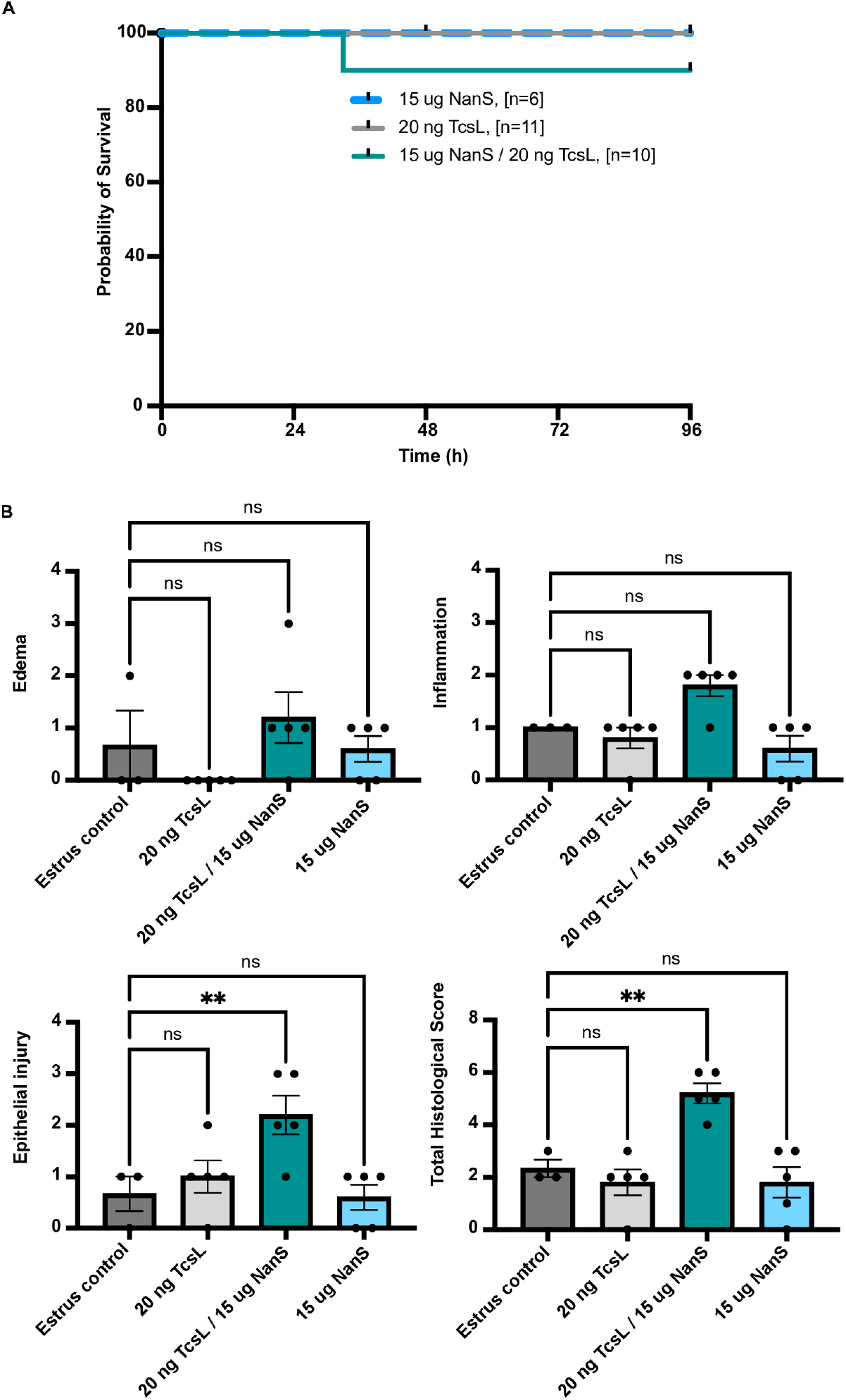
NanS works synergistically with TcsL to cause epithelial injury in estrus mice. **(A)** Survival curve of mice that were subcutaneously administered beta-estradiol on Day -2 to induce estrus, followed by transcervical instillation 15 μg NanS, 20 ng TcsL, or 15 μg NanS / 20 ng TcsL on Day 0. Animals were weighed/monitored for four days. Log-rank (Mantel-Cox) multiple comparison test was used with statistical significance set at a p value of <0.05. (**B**) Histological scoring of edema, acute inflammation, and epithelial injury of uterine tissues at end of study from estrus mice transcervically instilled with PBS-control, 20 ng TcsL alone, 20 ng TcsL / 15 μg NanS, or 15 μg NanS alone. Dunnett’s (one-way ANOVA) multiple comparisons test was used with statistical significance set at a p value of <0.05.

## Discussion

The *nanS* gene is present in all *P. sordellii* isolates [14], and yet the role of NanS in PSI remains elusive. We note that the *P. sordellii* ATCC9714 genome annotation contains a sialic acid catabolic operon suggesting that liberation of sialic acid could provide a nutrition benefit to the organism. In this study, we were particularly interested in roles NanS might have in enhancing cytotoxic effects. We expressed and purified a recombinant protein corresponding to the ATCC 9714 NanS sequence and found it to be a catalytically active sialidase (**Fig 1**). From the crystal structure, we were not able to discern any identifiable clues to suggest NanS operates differentially from other bacterial or viral sialidases.

We incubated recombinant NanS on HUVECs for 3 hours, and following removal, intoxicated the cells with recombinant TcsL for 24 hours. Our data indicate that NanS and TcsL work in a synergistic manner to increase cytotoxicity *in vitro* (**Fig 2**) but the effect was modest and required a high concentration of NanS. We next performed a cell rounding assay with Vero-GFP cells and found that NanS treatment of cells, followed by TcsL intoxication resulted in cells rounding at an increased rate compared to mock treated cells (**Fig 3**). This effect was evident at lower concentrations of TcsL and NanS and suggested that perhaps NanS is capable of cleaving sialic acids on the epithelium, most likely allowing for optimized TcsL-receptor access.

One *C. perfringens* study found that NanI was capable of increasing the binding of *C. perfringens* enterotoxin (CPE) and beta toxin to Caco-2 cells and HUVECs, respectively [10]. However, in the case of TcsL and its receptor SEMA6A, the interaction has been shown to occur in the absence of glycans, and de-glycosylation of SEMA6A did not alter TcsL/SEMA6A binding interactions [29,30]. Thus, we do not expect the sialidase activity of NanS to be directly affecting the interaction between TcsL and SEMA6A, but instead speculate that NanS is acting on glycosylation sites of the glycocalyx surrounding TcsL receptors. In addition, it is possible that NanS may be more impactful for TcsL binding and activity on differing cell types depending on the presence of adherent mucus, as was the case for *C. perfringens* NanI and CPE binding [31].

In observing the synergy in cell culture studies, we were motivated to explore the role of NanS *in vivo*. In our transcervical instillation model, TcsL alone can cause the major symptoms associated with *P. sordellii* infection when the mice are diestrus [28]. We therefore chose to instill 10 ng TcsL, an amount that resulted in approximately 50% survival by end of study at Day 4 in diestrus mice. By contrast, the co-instillation with NanS, caused all animals in diestrus to succumb to death before 24 hours, a result that was found to be statistically significant (**Fig 4A**). Since mucus is not present in the uterine horns of mice in diestrus (**S2 Fig A**), we speculate that NanS may work to degrade the sialoglycoprotein-rich glycocalyx atop the epithelium to aid TcsL in accessing its cell surface receptor.

To date in our lab, TcsL intoxication (up to 50 ng) in animals in estrus has not resulted in any signs of disease or sickness in the animals [28]. Vegetative PSI in animals in estrus resulted in significantly less death (30% mortality) compared to animals in diestrus (80% mortality) [28]. It is known, and we have observed, that there is a higher production of uterine mucus in estrus in contrast to diestrus (**S2 Fig B**). The mucus most likely plays a protective role for the host during pathogen exposure. In the case of *C. perfringens*, mucus has been shown to inhibit CPE, a major toxin responsible for severe human intestinal disease [31]. However, NanI, a sialidase produced by some *Clostridium perfringens* type F strains, was found to increase, *in vivo*, the action of CPE in the presence of mucus [31].

In PSI, we were curious if NanS might be responsible for break-down of sialic acid-containing mucus components in the uterus and increases *P. sordellii* pathogenesis in estrus mice. Perhaps, in estrus, TcsL requires NanS to break down mucus to allow the lethal toxin access to its receptor(s). Here, we found that 20 ng TcsL in an estrus mouse when co-instilled with NanS caused death in 10% of mice in the timeline of our experiment (**Fig 5A**). While the change in survival is not statistically significant, we noted significant changes in the tissue of the surviving mice, with the co-instilled animals experiencing increased inflammation and epithelial injury compared to control mice in estrus or mice instilled with 20 ng TcsL or 15 μg NanS alone. (**Fig 3B and 4B**). This experiment suggests a role for NanS in mucus and/or glycocalyx degradation in *P. sordellii* pathogenesis. Taken together, NanS could be considered an accessory virulence factor produced by *P. sordellii* to enhance TcsL-dependent mechanisms of *P. sordellii*-mediated disease.

## Methods and Materials

### Ethics statement

This study was approved by the Institutional Animal Care and Use Committee (IACUC) at Vanderbilt University Medical Center (VUMC) and performed using protocol M1700185-01. Our laboratory animal facility is AAALAC-accredited and adheres to guidelines described in the Guide for the Care and Use of Laboratory Animals. The health of the mice was monitored daily, and severely moribund animals were humanely euthanized by CO2 inhalation.

### Recombinant *P. sordellii* toxin purification

The *nanS* gene sequence was synthesized by GenScript and inserted into a pET-47b(+) vector under the T7 promoter using XmaI/XhoI restriction sites. Plasmid encoding His-tagged NanS (pBL974) was transformed into BL21Star *E. coli*, and two liters of LB medium (supplemented with 50 mg/liter kanamycin and 1% glucose) were inoculated with an overnight culture to an optical density at 600 nm (OD600) of ∼0.1. Cells were grown at 37°C and 220 rpm. Expression was induced with 0.5 mM IPTG once cells reached an OD600 of 0.5 to 0.6. After 4 h, the cells were centrifuged and resuspended in 20 mM HEPES (pH 8.0), 300 mM NaCl, and protease inhibitors. An EmulsiFlex C3 microfluidizer (Avestin) was used to generate lysates. Lysates were then centrifuged at 40,000 × g for 20 min. Supernatant containing NanS was run over a Ni-affinity column. The eluted protein fraction was dialyzed in 20 mM HEPES (pH 6.9), 100 mM NaCl.

TcsL was amplified from *P. sordellii* strain JGS6382 and inserted into a pCHis1622 vector (MobiTec) using BsrGI/KpnI restriction digestion sites in the vector, as reported previously [18]. Plasmid encoding Histagged TcsL (pBL552) was transformed into *Bacillus megaterium* according to the manufacturer’s protocol (MoBiTec). Six liters of LB medium supplemented with 10 mg/liter tetracycline were inoculated with an overnight culture to an OD600 of ∼0.1. Cells were grown at 37°C and 220 rpm. Expression was induced with 5 g/liter of D-xylose once cells reached an OD600 of 0.3 to 0.5. After 4 h, the cells were centrifuged and resuspended in 20 mM HEPES (pH 8.0), 500 mM NaCl, and protease inhibitors. An EmulsiFlex C3 microfluidizer (Avestin) was used to generate lysates. Lysates were then centrifuged at 40,000 × g for 20 min. Supernatant containing toxin was run over a Ni-affinity column. Further purification was performed using anion-exchange chromatography (HiTrap Q HP, GE Healthcare) and gel filtration chromatography in 20 mM HEPES (pH 6.9), 50 mM NaCl.

### Sialidase activity assays

For each sialidase activity assay, 20 μL of recombinant WT NanS was added to 60 μL of 100 mM sodium acetate buffer (pH 5.5) in a 96-well flat bottom plate. Then, 20 μL of 4 mM 5-bromo-4-chloro-3-indolyl α-D-N-acetylneuraminic acid (Sigma) was added, and the plate was incubated at 37°C. The absorbance at 620 nm was measured over time using a BioTek Cytation5 plate reader.

### X-ray crystallography of NanS

*P. sordellii* NanS was run over a size-exclusion column (Superdex-75) in 20 mM HEPES (pH 6.9), 50 mM NaCl buffer. The resulting sample was concentrated to 54 mg/ml and set up in broad matrix crystallization screens to identify crystal growth conditions. Final optimized crystallization conditions were 0.1 M Bis-Tris (pH 6.5), 0.2 M ammonium acetate, 25 % polyethylene glycol 3350, and crystals were harvested and cryoprotected in mother liquor with 20% glycerol. X-ray diffraction data were collected at the LS-CAT beamline (Advanced Photon Source, Argonne National Laboratory). Data were processed with autoPROC [32–36], and molecular replacement was performed in Phaser [37] with the structure of the sialidase from *Salmonella typhimurium* (Protein Data Bank code 1DIM) as search model to phase the data. Refinement was performed in Phenix [38], and Coot was used for model building [39]. Software was curated by SBGrid [40]. Coordinates and data have been deposited in the Protein Data Bank under accession code PDB ID 8T43.

### HUVEC culture and viability assays

Human umbilical vein endothelial cells (HUVECs) were maintained in Vascular Cell Basal Media supplemented with Endothelial Cell Growth Kit-VEGF, designed to keep a low serum culture environment. Cells were cultured at 37 °C with 5% CO2 and seeded into 96-well plates and allowed to grow until confluent (up to 2 days). For 3 hours, 10 μM NanS was incubated on cells, after which it was removed and replaced with cell media. TcsL or mock buffer was incubated on the cell for 24 hours, and ATP as a readout of viability was measured using the CellTiter-Glo luminescent cell viability assay (catalog number G7573; Promega).

### Vero-GFP cell culture and rounding assays

Vero-GFP cells were maintained in DMEM supplemented with 10% fetal bovine serum and cultured at 37 °C with 5% CO2. Cells were seeded into 96-well plates at 24,000 cells per well and allowed to grow overnight. For 3 hr, 100 nM NanS was incubated on cells after which it was removed and replaced with cell media. TcsL or mock buffer was incubated on the cell, and every 30 min for 20 hr, a GFP image was taken. From these images, the total number of rounded and non-rounded cells were counted.

### Animals and housing

All mouse experiments were approved by IACUC. C57BL/6J mice (all females, age 9 to 12 weeks) were purchased from Jackson Laboratories and were housed five to a cage in a pathogen-free room with clean bedding and free access to food and water. Mice had 12 hr cycles of light and dark.

### Virulence studies

*In vivo* virulence studies were conducted using a transcervical instillation model [28]. In brief, mice were anesthetized, and a speculum was inserted into the vaginal cavity to allow for dilation and passage of a flexible gel-loading pipette tip through the cervix and transfer of recombinant protein directly into the uterus. Following instillation, a cotton plug applicator was inserted into the vagina, and a cotton plug was expelled from the applicator and into the vaginal cavity using a blunt needle. Mice were monitored daily for morbidity and signs of sickness. Mice were humanely euthanized by CO2 inhalation when moribund or at end of study. In some cases, the uterus was harvested, fixed in Carnoy’s solution, paraffin-embedded, and processed for histology.

### Statistical analysis

Statistical testing and graphical representations of the data were performed using GraphPad Prism. Statistical significance was set at a P ≤ 0.05 for all analyses (*, P ≤ 0.05; **, P ≤ 0.01; ***, P ≤ 0.001; ****, P ≤ 0.0001). The Log-rank (Mantel-Cox) multiple comparison test was used for survival curve comparisons. Sidak’s multiple comparison (2-way ANOVA) test was used to compare two groups and Dunnett’s multiple comparison (2-way ANOVA) test was used to compare multiple groups. Ordinary one-way ANOVA test was used to compare histological scoring.

## Acknowledgements

We gratefully acknowledge Vanderbilt University Medical Center Translational Pathology Shared Resource for assistance with tissue embedding. The Translational Pathology Shared Resource is supported by NCI/NIH Cancer Center Support Grant P30CA068485.

## Author Contributions

**Conceptualization:** SCB, DBL

**Data curation:** SCB, HK

**Formal analysis:** SCB, HK, MKW, DBL

**Funding acquisition:** SCB, DBL

**Investigation:** SCB, HK

**Methodology:** SCB

**Project administration:** DBL

**Resources:** SCB

**Supervision:** DBL

**Validation:** SCB, HK

**Visualization:** SCB, HK

**Writing – original draft:** SCB

**Writing – review & editing:** SCB, DBL

## Supporting Information

**S1 Fig. ATCC 9714, compared to other *P. sordellii* strains, had an increased cytotoxicity and resulted in cell rounding following NanS treatment**. (**A**) Viability assay of Vero cells treated with 10 μM NanS for 3 hr followed by incubation of 48-hr growth supernatants (diluted 10-fold) of *P. sordellii* strains: ATCC 9714, DA108, HH-310, ORE2019. At 24 hr, ATP was measured as a readout of viability and normalized to signal from either NanS treated cells or mock treated cells. Assay was performed one time. (**B**) Vero cells treated with 48-hr supernatant from ATCC 9714 strain (left) and mock treated cells (right).

**S2 Fig. Luminal mucosa is not present in uteri of mice during diestrus but is present during estrus**. 20x magnification of Periodic Acid Schiff staining of uteri fixed in Carnoy’s solution of an animal in diestrus (**A**) and estrus (**B**).

